# Definitive benchmarking of DDA and DIA for host cell protein analysis on the Orbitrap Astral in a regulatory-aligned framework

**DOI:** 10.1101/2025.07.31.667876

**Authors:** Somar Khalil, Jenny T.C Ho, Michel Plisnier

**Affiliations:** GSK, Rue de l’Institut, 89, Rixensart, 1330, Belgium; ThermoFisher Scientific, Boundary Way, Hemel Hempstead, HP2 7GE, United Kingdom

**Keywords:** Astral, biopharmaceuticals, host cell proteins, label free, LC-MS/MS, mass spectrometry, orbitrap, proteomics

## Abstract

Host cell proteins (HCPs) are critical quality attributes in biotherapeutics and require accurate, protein-resolved quantification beyond the coverage limits of immunoassays. The Orbitrap Astral mass spectrometer was evaluated for label-free HCP analysis through a direct comparison of data-dependent acquisition (DDA, Top80) and data-independent acquisition (DIA, 4 m/z windows). Deterministic protein inference was implemented to ensure invariant protein grouping across datasets and software versions. Quantification was anchored using a whole-proteome stable isotope–labeled HCP standard, with identification error controlled through empirical false discovery proportion estimation. Both acquisition modes quantified total HCP content with high linearity (R^2^ > 0.99) and total error within ±30% acceptance limits across a seven-point spike-in series. DIA achieved greater analytical depth, identifying 45% more proteins and 68% more peptides than DDA, with substantially reduced missingness. Protein-level fold-change behavior was evaluated using hierarchical Bayesian regression, with slopes centered near unity for DIA and systematic compression observed for DDA. Abundance-stratified resampling analysis demonstrated trueness and precision across the dynamic range and defined lower limits of quantification of approximately 0.6 ppm for DIA and 1.6 ppm for DDA. Both acquisition strategies accurately report global HCP content, while DIA provides improved coverage, fold-change fidelity, and sensitivity for low-abundance impurities. This study demonstrates that untargeted MS-based HCP measurements can be analytically qualified for use in regulated biopharmaceutical settings.

## 1. Introduction

Host cell proteins (HCPs) are process-related impurities originating from expression systems used in recombinant biotherapeutic production [1,2]. As critical quality attributes, residual HCPs require quantitative monitoring because they can compromise product stability, contribute to immunogenicity, or lead to adverse clinical outcomes [3–6]. HCP monitoring has traditionally relied on enzyme-linked immunosorbent assays, which offer high throughput but lack protein-level resolution and depend on polyclonal antibody reagents [7,8]. These limitations have motivated the use of liquid chromatography–tandem mass spectrometry (LC–MS/MS) as an orthogonal platform capable of direct, protein-resolved HCP measurement [9,10].

Among LC–MS/MS acquisition strategies, data-dependent acquisition (DDA) remains widely used but is affected by stochastic precursor selection, leading to missing values and reduced reproducibility for low-abundance species [11]. Data-independent acquisition (DIA) systematically fragments all precursors within defined m/z windows, improving data completeness and reproducibility at the expense of increased spectral complexity [12–14]. Recent instrument developments have reduced this trade-off. The Orbitrap Astral MS couples high-resolution Orbitrap MS1 detection with an ultra-fast Asymmetric Track Lossless (Astral) MS2 analyzer. This architecture enables true parallel acquisition with scan rates exceeding 200 Hz, narrow DIA isolation windows (2–4 Th), and high ion utilization [15,16]. These characteristics are particularly relevant for HCP analysis, where sensitivity and dynamic range are constrained by a dominant product matrix [17–19].

Despite these advances, adoption of DIA in regulated workflows has been limited by concerns related to computational deconvolution and statistical error control. While target–decoy strategies are well established for DDA, some DIA pipelines require careful parameterization to achieve comparable stringency [20–22]. As regulatory expectations increasingly emphasize empirical error characterization, narrow-window, high-resolution DIA benefits from orthogonal validation such as entrapment-based estimation of the false discovery proportion (FDP) to support analytical specificity [23–29].

A further challenge in MS-based HCP analysis is the absence of ground truth for endogenous impurities. While stable isotope–labeled (SIL) peptides are often used, they do not reproduce the structural and physicochemical diversity of intact HCPs [30–32]. Spiked-in heterologous whole-protein mixtures provide broader coverage and enable evaluation of linearity, accuracy, and precision in line with ICH Q2(R2) guidelines [32] but may not fully reflect biological heterogeneity or matrix interactions of endogenous impurities. Whole-protein SIL-HCP mixtures derived from the production cell line provide a more representative alternative. When spiked into a purified drug substance matrix, they mimic the complexity of native process-related impurities and enable performance evaluation under realistic conditions [33,34]. Their diversity supports worst-case impurity behavior and allows traceable benchmarking across the analytical range for assessing trueness, precision, and recovery per ICH Q2(R2). Used routinely, SIL-HCP standards can also function as system suitability materials, verifying recovery and bias against predefined acceptance limits [35,36].

As biopharmaceutical development shifts toward risk-based impurity control, there is a need to move beyond proteome coverage metrics toward qualification of MS-based workflows. Here, we benchmark the Orbitrap Astral MS for label-free HCP quantification by directly comparing an optimized DDA method (Top80) with a narrow-window DIA method (4 m/z non-staggered). Performance was evaluated using a Chinese hamster ovary (CHO)–derived SIL-HCP standard spiked into a NISTmAb matrix, empirical false discovery proportion (FDP) estimation using complementary entrapment strategies, deterministic protein inference via greedy parsimony constrained by Occam’s razor [37,38], and abundance-stratified assessment of linearity, accuracy, and limits of quantification across a seven-point concentration series.

## 2. Experimental Section

### 2.1 Sample preparation

NISTmAb reference material (humanized IgG1κ, NIST RM 8671) was used as the model product matrix. A SIL-HCP standard derived from CHO cells (Sigma) was spiked into the mAb at seven levels (L1–L7: 4%, 6%, 8%, 10%, 12%, 14%, and 16% w/w relative to the mAb). Sample preparation was adapted from a previous study with minor modifications [34]. Each replicate containing 20 µg of mAb protein was solubilized in 0.1% RapiGest (Waters) and 50 mM triethylammonium bicarbonate (Thermo Fisher Scientific). Proteins were reduced with 8 mM dithiothreitol at 50 °C for 45 min and alkylated with 16 mM iodoacetamide in the dark at room temperature for 30 min. Digestion was performed overnight at 37 °C using a Trypsin/Lys-C mix (Thermo Fisher Scientific) at a 1:40 enzyme-to-substrate ratio. The reaction was quenched with 0.5% trifluoroacetic acid (Biosolve). Peptides were desalted using C18 spin columns (Thermo Fisher Scientific), dried by vacuum centrifugation, and reconstituted in 0.1% formic acid (Biosolve). Each digest was spiked with four MassPREP protein digest standards (P02769, P00330, P00489, and P00924; Waters) at 0.37, 1.02, 1.37, and 5.14 fmol/µL, respectively. All samples were analyzed in technical triplicate by DDA and DIA on the Orbitrap Astral MS.

### 2.2 Liquid chromatography-mass spectrometry analysis

Tryptic peptides corresponding to a 500-ng theoretical mAb load per injection were analyzed using a Vanquish Neo UHPLC system (Thermo Fisher Scientific) operated in direct injection mode. Peptides were separated on an EASY-Spray PepMap Neo C18 column (100 Å, 2 µm, 75 µm ID x 50 cm, Thermo Fisher Scientific) maintained at 55 °C. Mobile phase A consisted of 0.1% formic acid in water, and mobile phase B consisted of 80% acetonitrile, 20% water, and 0.1% formic acid. The LC gradient was as follows: 3–8% B over 3 min at 500 nL/min; 8–32% B over 22 min at 300 nL/min; 32–50% B over 5 min; ramp to 99% B over 1 min at 500 nL/min; and hold at 99% B for 4 min. Electrospray ionization was performed at 1.8 kV. The ion transfer capillary was set to 285 °C, and the ion funnel RF level to 40. For DDA, MS1 scans were acquired in the Orbitrap at 240K resolution (at m/z 200) over an m/z range of 380–1500, with a maximum injection time (maxIT) of 20 ms and automatic gain control (AGC) target of 5 × 10^6^. A Top80 method selected precursors with charge states ≥2 and intensities ≥5 x 10^3^, using a quadrupole isolation width of 0.7 Th. MS2 spectra were acquired in the Astral analyzer using higher-energy collisional dissociation (HCD) at 27% normalized energy, with an m/z range of 120–2000, maxIT of 6 ms, and AGC target of 5 × 10^3^. For DIA, MS1 scans were also acquired at 240K resolution over m/z 380–980, with a 10 ms maxIT and AGC target of 5 × 10^6^. MS2 spectra were acquired in the Astral analyzer using non-overlapping 4 Th precursor isolation windows. Fragment ions were measured over m/z 150–2000 using HCD at 25%, 4 ms maxIT, and AGC target of 5 × 10^4^.

### 2.3 Data processing and protein inference

Raw data were processed using vendor-neutral software: SpectroMine (v5.0, Biognosys) for DDA and Spectronaut (v20.0, Biognosys) for DIA, with the latter operated in DirectDIA library-free mode. Both searches were conducted using identical parameters against a composite FASTA database comprising 83419 Cricetulus griseus entries from UniProt (Swiss-Prot and TrEMBL), the NISTmAb heavy and light chain sequences, and the four MassPREP digest standards. Trypsin/P was specified as the proteolytic enzyme with up to one missed cleavage.

Mass tolerances were set to automatic. Fixed modifications included carbamidomethylation (C) and stable isotope labels on arginine (+10 Da) and lysine (+8 Da), while variable modifications included methionine oxidation and N-terminal acetylation. For targeted searches of the MassPREP standards, carbamidomethylation was specified as the sole fixed modification. Peptides were filtered to lengths of 7–30 amino acids and precursor charge states between 2^+^ and 4^+^. A uniform q-value threshold of 0.01 was applied at the precursor, peptide, and protein levels. In Spectronaut, confidence filtering used q-value and posterior error probability thresholds of 0.01 at both run and experiment levels [39]. No normalization or imputation was applied at this stage. Quantification was performed at the MS1 level for DDA using extracted ion chromatograms and at the fragment-ion level for DIA using summed peak areas.

Protein inference was performed independently of the search engines using a deterministic pipeline. Following 1% FDR filtering, stripped peptide sequences were re-annotated against an *in silico* tryptic digest of the *C. griseus* proteome. Protein groups were constructed using a greedy parsimony algorithm, followed by Occam’s razor filtering requiring at least one unique peptide per group. For each group, the lead protein was defined as the member supported by the highest number of constituent peptides. Groups without unique peptide evidence were retained without further subdivision. All computational workflows were executed on a virtual machine equipped with a 64-core AMD EPYC processor and 256 GB RAM.

Downstream statistical analysis and visualization were performed in Python (v3.12) using in-house scripts based on NumPy, pandas, and seaborn. Bayesian modeling was implemented in PyMC, with posterior diagnostics evaluated using ArviZ. Final figures were assembled in GraphPad Prism (v10.5).

### 2.4 Empirical FDP estimation

Empirical estimation of FDP was performed using a search-centric entrapment strategy adapted from Wen et al. [22]. Target peptides were defined from a reference search against the unmodified database. Two entrapment databases were constructed and analyzed separately. In the first approach, a shuffled peptide database was generated by randomly permuting internal amino acid sequences of identified *C. griseus* peptides while preserving the C-terminal residue (K/R). Peptides identical to native sequences were removed, and the final set was down-sampled to maintain a 1:1 ratio of target to entrapment peptides. In the second approach, a trimmed foreign proteome database was generated from an *in silico* tryptic digest of Arabidopsis thaliana. Only peptides with no sequence overlap with the *C. griseus* peptidome were retained, and the resulting set was down-sampled to match the size of the target peptide space. Each entrapment FASTA was concatenated with the target database and searched under identical conditions. FDP was estimated at multiple q-value thresholds (*τ*) using two estimators.

- The combined FDP estimator was defined as:

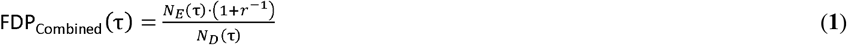

where *N*_*E*_ (*τ*) is the number of identified entrapment peptides below threshold (*τ*), *N*_*D*_ (*τ*) is the total number of discoveries, and *r* is the target to entrapment ratio in the database (*r* = 1).

- The paired FDP estimator was calculated as:

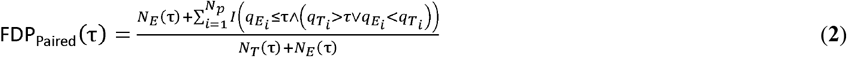

where each of the *N*_*p*_ target-entrapment peptide pairs (*T*_*i*_, *E*_*i*_) was evaluated using the indicator function I, which counts entrapments identifications occurring alone or with a lower q-value than their corresponding targets. *N*_*T*_ (*τ*) and *N*_*E*_ (*τ*) denote the numbers of target and entrapment peptides identified below threshold (*τ*), respectively.

### 2.5 Statistical analysis

#### 2.5.1 Data filtering and normalization

Post-inference, peptide-level data were processed using a sequential quality control pipeline. At each spike-in level, outlier peptides were identified using the modified Z-score approach, with peptides satisfying |*Zi*| > 2.6 were excluded. The score was calculated as:

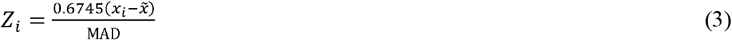

where *x*_*i*_ is the log_2_-transformed intensity, 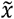 is the median intensity, and MAD is the median absolute deviation.

Intra-protein consistency was applied by removing peptides whose median log_2_ intensity deviated by more than 30-fold from the median intensity of the corresponding protein. Data were corrected for loading variation using total sum scaling applied within each spike-in level. A precision filter was subsequently applied, excluding peptides with a coefficient of variation (CV%) greater than 25% across technical triplicates. log_2_ intensity density distributions before and after filtering were compared to assess the impact of the quality control steps (**Supplementary Figure S1**).

#### 2.5.2 Hierarchical Bayesian modeling of differential linearity

Protein-level fold-change (FC) behavior was modeled using a hierarchical Bayesian regression with partial pooling. Individual protein linearity parameters (β_j_) were modeled as draws from a common group-level distribution. Observed log_2_FC values (y_ij_) were modeled as a function of the expected log_2_FC (*x*_*i*_) across spike-in levels using a Student’s t likelihood. The model was specified as:

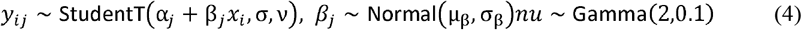

where µ_*β*_ represents the group-level mean slope, and the degrees-of-freedom parameter (ν) is estimated from the data using a weakly informative Gamma hyperprior. Posterior inference was performed using the No-U-Turn Sampler [40]. Model diagnostics included evaluation of the Gelman–Rubin statistic 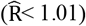 and inspection of trace plots.

#### 2.5.3 Global HCP accuracy profile

Accuracy of total HCP quantification was evaluated using accuracy profiles based on the total-error approach. Trueness was defined as the relative bias from the nominal spike level, and precision was expressed using two-sided tolerance intervals. For each spike level (*j)* with (*n*_*j*_) replicate measurements (*Y*_*ij*_) of the nominal concentration (*µ*_*j*_), relative bias was calculated as:

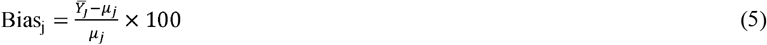

With

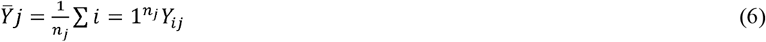

The within-level standard deviation was estimated as:

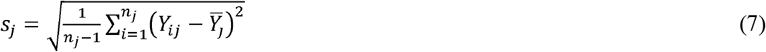

Two-sided tolerance intervals at level j were defined as:

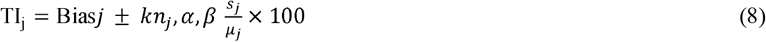

Two forms of tolerance factors were considered. β-expectation tolerance intervals were calculated assuming normally distributed data with unknown variance, using:

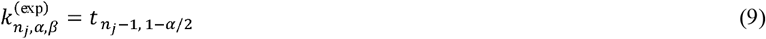

where (*t*_*v,p*_) denotes the *p*-quantile of the Student-t distribution with *v* = *n*_*j*_ − 1 degrees of freedom. For *n*_*j*_ = 3, *k* ≈ 4.303.

In addition, 95%/95% content tolerance intervals were computed using Hahn–Meeker tolerance factors [41], with *k* = 9.916 for *n*_*j*_ = 3:

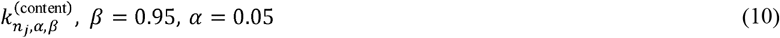

A predefined acceptance criterion of ±30% total error was applied across all spike levels.

#### 2.5.4 Stratified bootstrap analysis

Quantification fidelity across the abundance range was evaluated using a stratified non-parametric bootstrap. At each spike-in level, proteins were stratified into four abundance percentiles based on concentration (ppm): P0–15, P15–35, P35–50, and P50–100. Within each stratum, protein abundances were resampled with replacement (12K iterations) to generate bootstrap distributions of the stratum mean. Point estimates were defined as the mean of the bootstrap distribution, and uncertainty was quantified using two-sided 95% percentile confidence intervals (CI). The estimand was the stratum mean, aligning the accuracy assessment with the level of inference. Estimates were normalized within each stratum to a fixed reference derived at the L4 spike level, computed once by bootstrap and treated as constant to avoid variance propagation.

Accuracy was evaluated by comparing the normalized stratum mean bias and its CI against a predefined ±35% total-error acceptance limit. Linearity within each stratum was assessed by regression of stratum mean estimates against nominal concentrations, with a coefficient of determination threshold of R^2^ ≥ 0.98. The lower limit of quantification (LLOQ) was defined as the lowest abundance stratum satisfying both the accuracy and linearity criteria.

## 3. Results and Discussion

### 3.1 Acquisition-dependent sampling and data completeness

A head-to-head comparison on the Orbitrap Astral revealed clear differences between DIA and DDA in proteome coverage and data completeness across all spike-in levels. DIA identified more protein groups and peptides than DDA at every concentration (**Figure 1A–B**) and yielded a higher average number of peptides per protein (**Figure 1C**). DDA produced a slightly higher absolute number of single-hit proteins, while the proportion of single-hit identifications relative to total protein groups was consistently lower for DIA across all spike levels (**Figure 1D**). Differences were observed in data completeness. At lower spike levels, DDA showed pronounced peptide missingness, with more than 50% of peptides absent at the lowest concentrations, consistent with intensity-driven precursor selection (**Figure 1E**) [42-45]. In contrast, DIA maintained high peptide detection rates across the range. Overlap analysis showed that 87% of proteins and 96% of peptides identified by DDA were also detected by DIA, while DIA uniquely identified a large number of additional features (**Figure 1F–G**).

**Figure 1.**
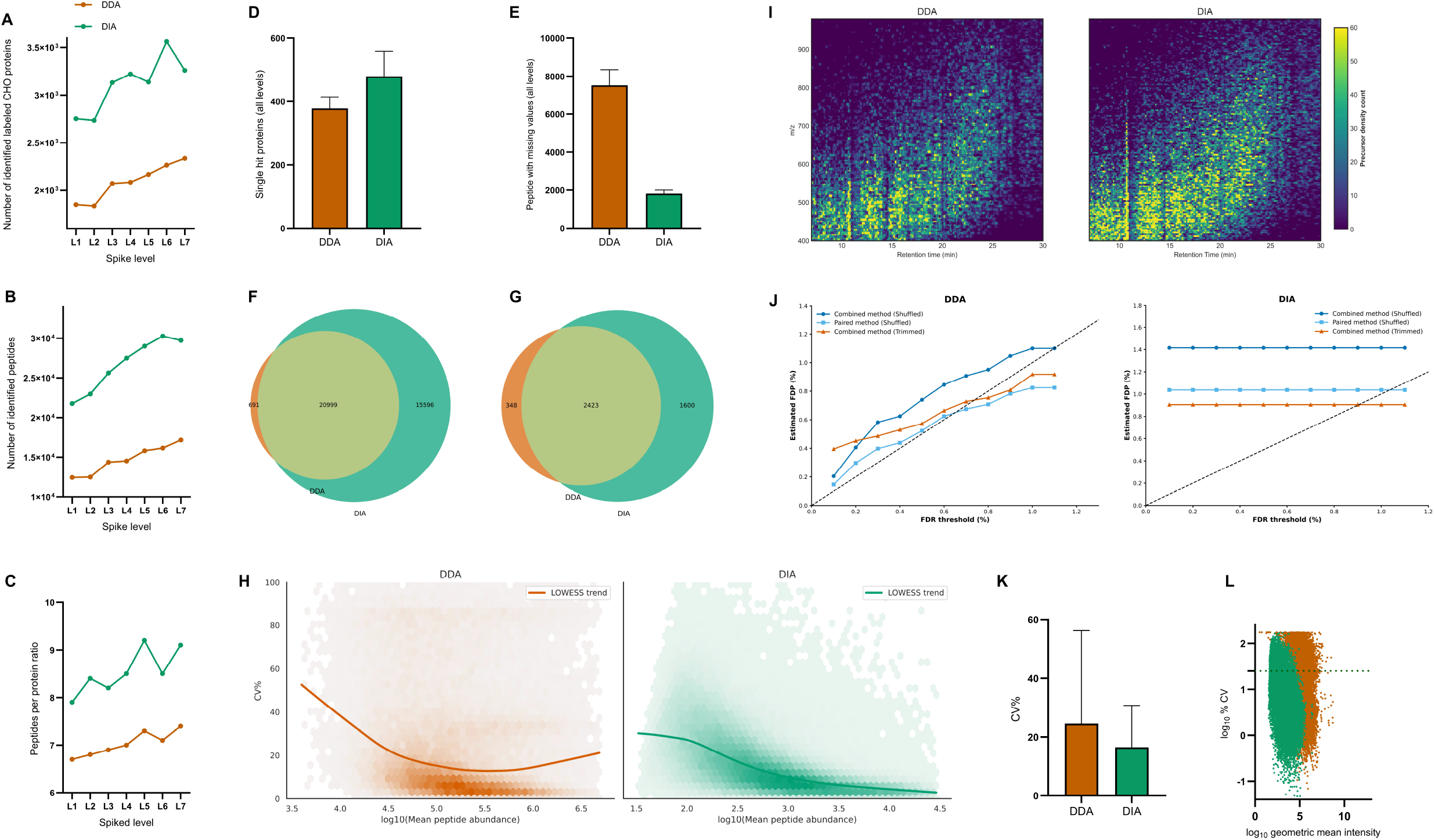
Comparative evaluation of DDA and DIA workflows for proteome coverage, data completeness, and quantitative precision. Line plots show the number of identified protein groups (**A**), peptides (**B**), and the average number of peptides per protein (**C**). Bar charts compare the number of single-hit proteins (**D**) and peptides with missing values (**E**). Venn diagrams summarize overlap in identified proteins (**F**) and peptides (**G**). Hexbin plots show peptide-level CV% as a function of abundance with LOWESS smoothing (**H**). Two-dimensional density maps illustrate precursor sampling across retention time and m/z space (**I**). Empirical FDP estimates derived from paired and combined entrapment strategies are shown in (J). Mean peptide-level CV% across all concentrations is shown in (**K**). Scatter plots of log-transformed CV% versus log-transformed geometric mean intensity are shown in (**L**). Error bars represent standard deviation across spike-in levels.

Acquisition-specific sampling patterns were evident when precursor distributions were visualized across retention time and m/z space (**Figure 1I**). DDA showed sparse and banded sampling, whereas DIA showed uniform and continuous coverage. These differences were reflected in the peptide-level CV% distributions, which were lower and narrower for DIA across all spike-in levels (**Figure 1K–L**). Peptide-level CV% trends were characterized using LOWESS smoothing, revealing stable variability for DIA across intensities, while DDA showed increased dispersion at both low and high abundance levels (**Figure 1H**).

### 3.2 Empirical estimation of precursor-level FDP by entrapment

Empirical FDP analysis showed effective statistical error control for both acquisition modes, with estimator-dependent behavior. In the DDA workflow, all three estimators (paired-shuffled, combined-shuffled, and combined-trimmed) returned FDP values below the nominal 1% threshold across precursor-level q-value cutoffs between 0 and 1% (**Figure 1J, left**). FDP curves showed close agreement across estimators, with a shallow non-monotonic dip observed for the combined-shuffled estimator at higher q-value cutoffs [46,47]. In the DIA workflow, FDP estimates remained invariant, reflecting a strongly bimodal q-value distribution. At the 1% operating point, the paired-shuffled and combined-trimmed estimators remained at or below the nominal FDP, while the combined-shuffled estimator yielded an elevated estimate of approximately 1.5% (**Figure 1J, right**). This divergence reflects known behavior of shuffled-sequence decoys in highly multiplexed DIA data, where increased susceptibility to spurious matches inflates entrapment counts. Agreement between the paired-shuffled and combined-trimmed estimators supports effective precursor-level error control at the nominal 1% threshold. While empirical underestimation of error rates has been reported for some DIA pipelines, the configuration and filtering applied here yielded FDP estimates aligned with the intended level of statistical control [22,48,49].

### 3.3 Hierarchical modeling reveals acquisition-mode effects on protein-level fold-change

Protein-level quantification used the Hi3 label-free approach, with abundances computed from the summed intensities of the three most intense quantifiable peptides per spike-in level [34,50]. Across all concentrations, DIA quantified more peptides and proteins than DDA (**Figure 2A–B**). Overlap analysis showed a shared core of approximately 1509 quantifiable proteins, with DIA uniquely quantifying an additional ∼1600 proteins (**Figure 2C**). Quantitative agreement between acquisition modes was examined using Bland–Altman analysis of log_2_FCs across representative spike-in comparisons (**Figure 2D**). Mean bias between DDA and DIA was close to zero; however, the 95% limits of agreement spanned approximately ±2 log units, corresponding to up to four-fold differences at the individual protein level. This dispersion indicates that, despite similar global trends, protein-level FC estimates are not interchangeable between acquisition modes.

**Figure 2.**
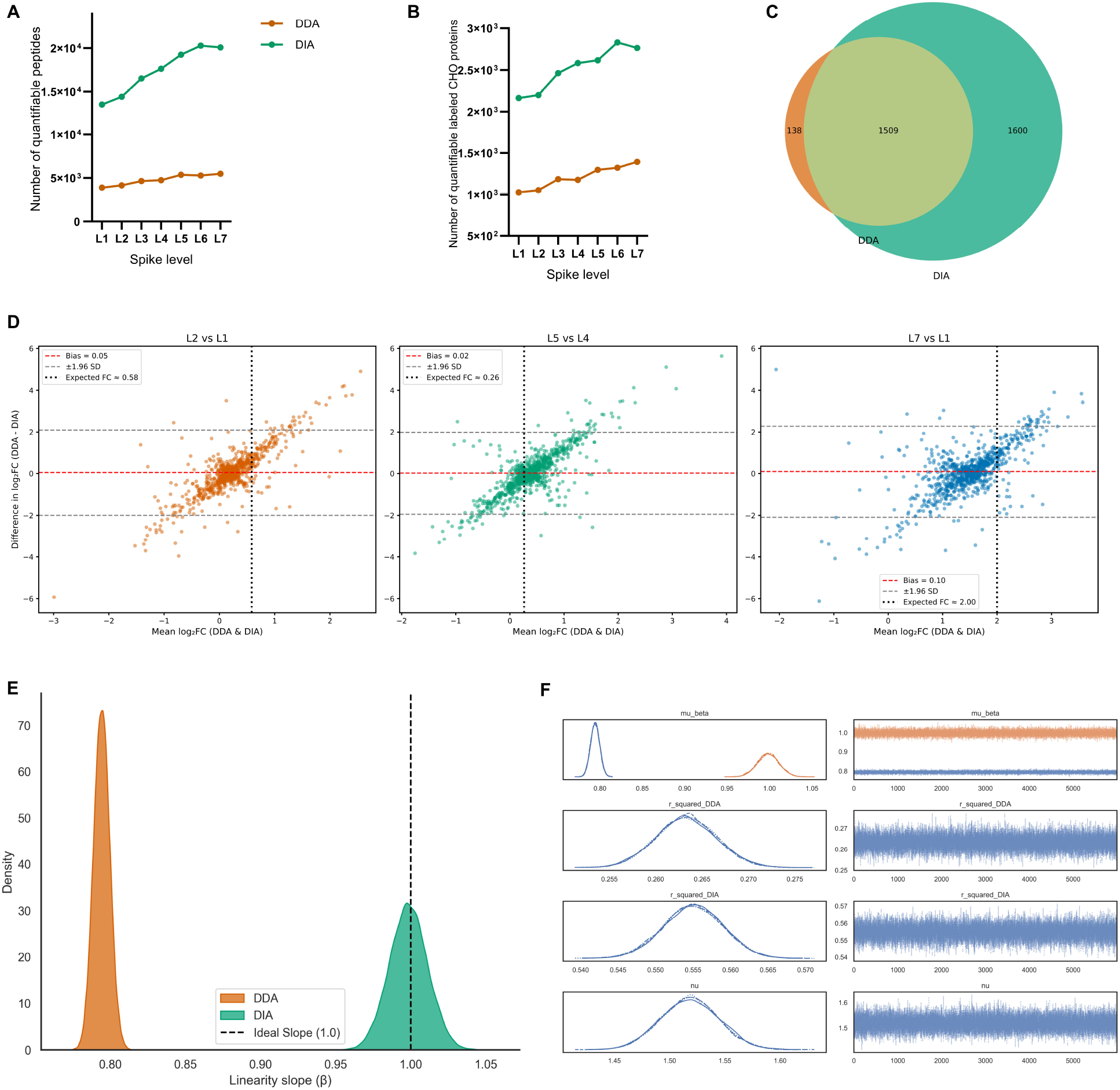
Protein-level quantification and FC behavior for DDA and DIA. Line plots show the number of quantifiable peptides (**A**) and proteins (**B**) across seven spike-in levels. (**C**) Venn diagram summarizing overlap in quantifiable proteins between acquisition modes. (**D**) Bland– Altman plots comparing log_2_FC between DDA and DIA for representative concentration contrasts, with mean bias (solid line) and 95% limits of agreement (dashed lines). (**E**) Posterior density distributions of the group-level linearity slope parameter (µ_β_) from hierarchical Bayesian regression, shown relative to the ideal slope of 1.0. (**F**) MCMC diagnostics including posterior densities (**left**) and trace plots (**right**) for key model parameters.

FC linearity was assessed using a hierarchical Bayesian regression model. Posterior distributions of the group-level slope parameter (µ_β_) were centered near unity for DIA, whereas DDA showed systematic slope compression with µ_β_ ≈ 0.8 (**Figure 2E**). Markov chain Monte Carlo diagnostics reported stable sampling behavior for all parameters (**Figure 2F**).

### 3.4 Total HCP abundance and total-error performance

Absolute HCP abundance was determined by calibrating summed top-three peptide intensities of the MassPREP standards against known molar inputs. The resulting response factor was used to convert aggregated HCP signal to mass (ng) by molecular-weight scaling. Measured and theoretical HCP abundances were proportional for both acquisition modes, with R^2^ exceeding 0.99 across the evaluated range (**Figure 3A–B**).

**Figure 3.**
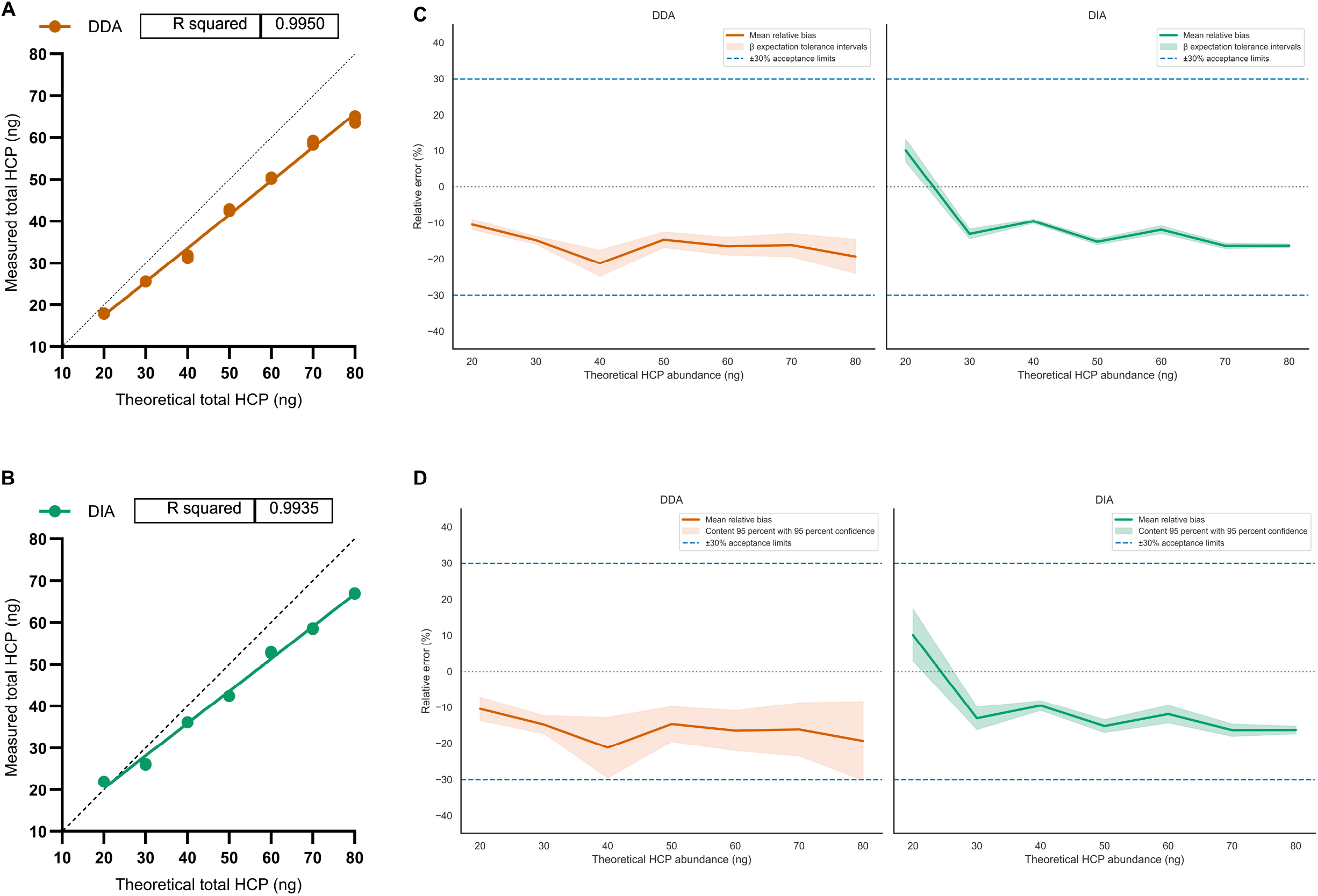
Measured versus theoretical total HCP abundance for DDA (**A**) and DIA (**B**), with linear regression fits (solid lines) and the identity line (dashed). (**C**) Accuracy profiles based on the total-error approach using 95% β-expectation tolerance intervals (α = 0.05). Solid lines indicate mean relative bias; shaded regions denote tolerance intervals; dashed lines indicate ±30% acceptance limits. (**D**) Accuracy profiles using 95%/95% content tolerance intervals.

Accuracy profiles based on the total-error approach revealed mode-dependent bias patterns across the 20–80 ng range. DDA exhibited a negative bias ranging from approximately −10% to −20%, whereas DIA showed a concentration-dependent bias, with positive bias at the lowest level (∼+10%) and a shift toward −15% at higher concentrations (**Figure 3C**). For both workflows, the 95% β-expectation tolerance intervals (α = 0.05) remained within the predefined ±30% acceptance limits. The corresponding 95%/95% content tolerance intervals were wider, reflecting the limited number of technical replicates (n = 3), but likewise remained within acceptance bounds across all spike levels (**Figure 3D**).

### 3.5 Abundance-stratified accuracy profiling and practical LLOQ estimation

Proteins were stratified into abundance percentiles (P0–15, P15–35, P35–50, P50–100) using boundaries defined on the combined dataset to ensure comparability across acquisition modes. At each spike-in level, protein abundances within each stratum were resampled with replacement to generate distributions of the stratum mean. Accuracy profiles showed that both DDA and DIA maintained mean relative bias within the predefined ±35% acceptance limits across all strata and spike levels (**Figure 4A–B**). CIs were narrower for DIA, with the largest differences observed in the lowest abundance strata. Kernel density estimates of the stratum mean distributions were unimodal and shifted with increasing spike levels for both workflows (**Figure 4C–D**). Regression of normalized stratum means against normalized nominal abundances resulted in R^2^ ≥ 0.98 for all strata (**Figure 4E**). Applying combined criteria of accuracy (mean bias and CI within ±35%) and linearity (R^2^ ≥ 0.98), the practical LLOQ was estimated at approximately 1.6 ppm for DDA and 0.6 ppm for DIA, with DIA showing greater stability in the lowest abundance strata.

**Figure 4.**
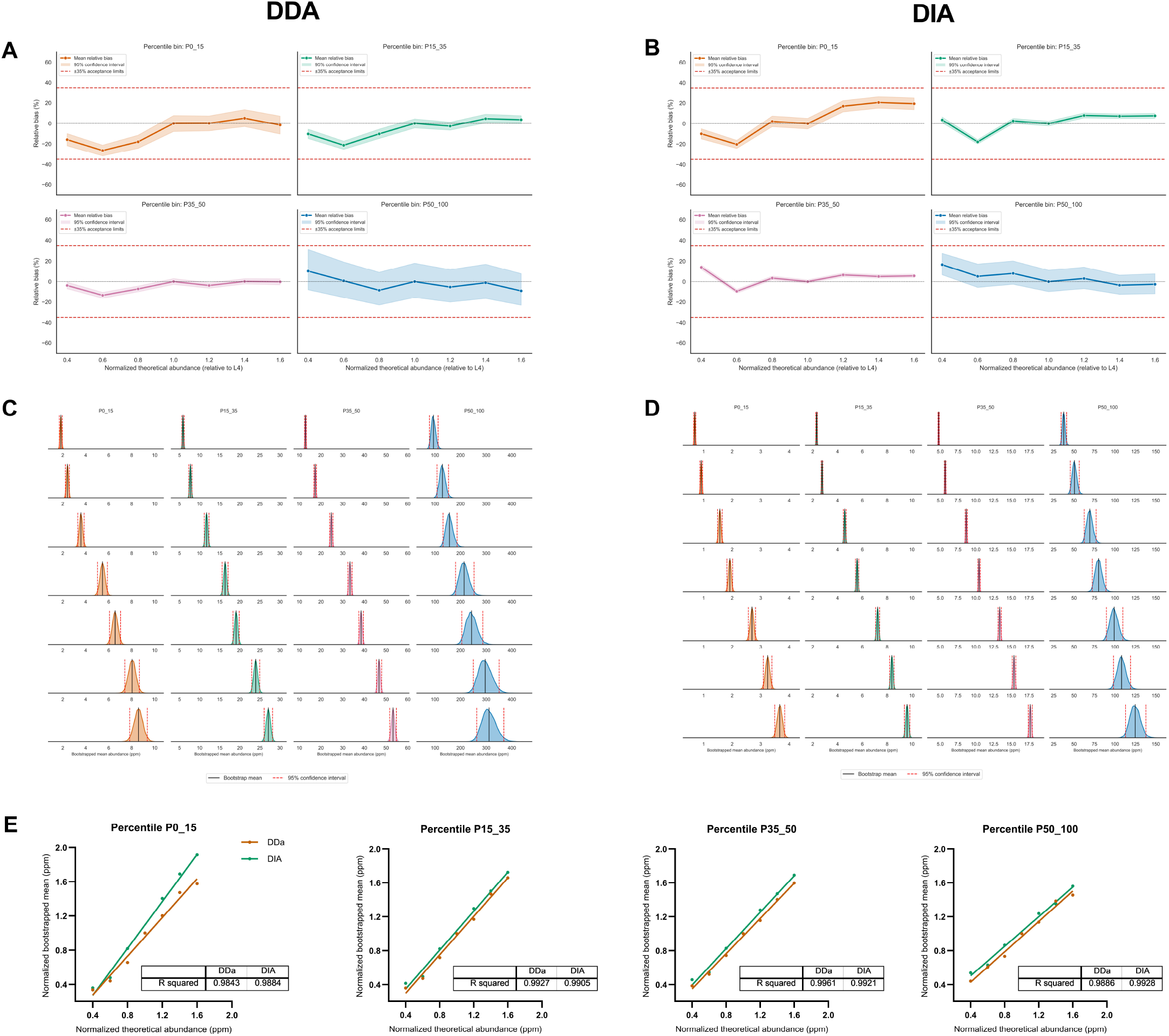
Accuracy profiles for DDA (**A**) and DIA (**B**) across abundance strata, showing mean relative bias (solid), 95% bootstrap CIs (shaded), and the ±35% limits (dashed). (C–D) Kernel density estimates of bootstrapped stratum means for DDA (**C**) and DIA (**D**) across spike-in levels (rows) and abundance bins (columns). (**E**) Linearity of normalized bootstrap means versus normalized nominal abundance for each stratum, with R2 values for both modes.

While global HCP accuracy was evaluated using β-expectation and 95%/95% tolerance intervals to support predictive assessment of assay performance, abundance-stratified accuracy was evaluated using bootstrap CIs to quantify uncertainty in stratum-level mean bias without distributional assumptions.

### 3.6 High-risk HCPs are quantified at the lowest spike level

At the lowest spike level (L1, 20 ng), both DDA and DIA quantified 19 of the 22 high-risk HCPs listed in **Table 1** [51]. Several well-characterized impurities showed close quantitative agreement between acquisition modes, including Cathepsin B (90 ppm in DDA vs 79 ppm in DIA) and Clusterin (74 ppm in both modes). Substantial mode-dependent differences were observed for a subset of proteins. Peroxiredoxin-1 was quantified at 103 ppm in DDA and 2 ppm in DIA, while Heat shock cognate 71 kDa protein was measured at 382 ppm in DDA and 119 ppm in DIA. Three high-risk proteins, Carboxypeptidase D, Glutathione S-transferase, and Transforming growth factor-β1, were not detected by either workflow at this concentration.

**Table 1.** High-risk HCPs quantified by DDA and DIA at the lowest spike level (L1: 20 ng). Reported values correspond to estimated protein abundance (ppm). ND, not detected.

Across the high-risk panel and the broader CHO proteome, DIA yielded higher median per-protein linearity, with a larger fraction of proteins exceeding an R^2^ ≥ 0.95 threshold compared with DDA (**Supplementary Figures S3 and S4**).

### 3.7 Framework for empirical qualification of untargeted MS-based HCP workflows

Qualification of untargeted MS-based HCP workflows can be structured around control of identification error, reproducible protein inference, and definition of the analytical range. In this study, these requirements were addressed through empirical FDP estimation, abundance-resolved accuracy assessment, and SIL-HCP–based calibration. Precursor-level specificity was evaluated using two independent entrapment strategies. Linearity of the calibrated response was assessed across a seven-point spike-in series, yielding R^2^ ≥ 0.99 for both acquisition modes. Trueness and precision were quantified using total-error accuracy profiles. For global HCP abundance, β-expectation values and 95%/95% content tolerance intervals were applied. For abundance-stratified analyses, non-parametric bootstrap confidence intervals were constructed at the stratum mean level. Both DDA and DIA met predefined acceptance limits of ±30% total error for global HCP abundance and ±35% within abundance-stratified bins. Integration of accuracy and linearity criteria yielded practical LLOQs of approximately 1.6 ppm for DDA and 0.6 ppm for DIA. Protein inference was implemented using a deterministic parsimony-based algorithm to ensure invariant grouping across datasets and software versions.

During method qualification, hierarchical modeling and stratified bootstrapping were used to characterize method performance. For routine application, a single SIL-HCP spike level is sufficient for benchmarking and system suitability assessment through verification of recovery and bias under conditions representative of endogenous impurities, without the full statistical workflow.

## 4. Conclusions

DDA and DIA acquisition on the Orbitrap Astral were evaluated for label-free HCP quantification. DIA delivered higher proteome coverage, tighter quantitative precision, improved fold-change stability across the measured dynamic range, and lower LLOQs, while both acquisition modes quantified HCP abundance within predefined acceptance limits at both aggregate and abundance-stratified levels. The analysis relied on whole-proteome SIL-HCP spiking, empirical FDP estimation, deterministic protein inference, and abundance-stratified performance assessment. These elements were sufficient to characterize specificity, linearity, trueness, precision, and analytical range for an untargeted MS-based HCP workflow.

The evaluation was limited to technical triplicates, a single monoclonal antibody matrix, and the CHO proteome. Reported LLOQs reflect standard tryptic digestion without product depletion or enrichment and are specific to the Orbitrap Astral instrument architecture. Performance characteristics may differ for alternative expression systems, product formats, or sample-preparation strategies. Generalizability of these performance characteristics is limited by the evaluated design. Extension to additional biologic modalities, host expression systems, or sample-preparation strategies requires re-estimation of quantitative bias and abundance-dependent precision under each condition. Instrument-specific effects related to scan speed and duty cycle similarly require empirical verification on alternative MS architectures.

Untargeted MS-based HCP analysis can be analytically qualified through empirical control of identification error, deterministic protein inference, and abundance-resolved performance characterization.

## Acknowledgments

The authors gratefully acknowledge Pascal Bourguignon and Nora Zaïm for their contributions to the laboratory work that supported this study. We also thank Jean-François Dierick for his valuable input and critical guidance on topics related to analytical method validation.

## Data availability

Raw LC–MS/MS data and SpectroMine/Spectronaut search results have been deposited with the ProteomeXchange Consortium via the MassIVE repository under identifier MSV000098998 (ProteomeXchange accession PXD067958; DOI:10.25345/C5T14V28X). Processed quantification tables and Python scripts are available at Zenodo (DOI:10.5281/zenodo.16883906).

## CRediT AUTHOR statement

**Somar Khalil:** Conceptualization, Methodology, Formal analysis, Investigation, Data Curation, Writing - Original Draft, Writing - Review & Editing. **Jenny T.C Ho**: Formal analysis, Review & Editing. **Michel Plisnier:** Writing - Review & Editing, Supervision, Project administration.

