## Supplementary for "Definitive benchmarking of DDA and DIA for host cell protein analysis on the Orbitrap Astral in a regulatory-aligned framework"

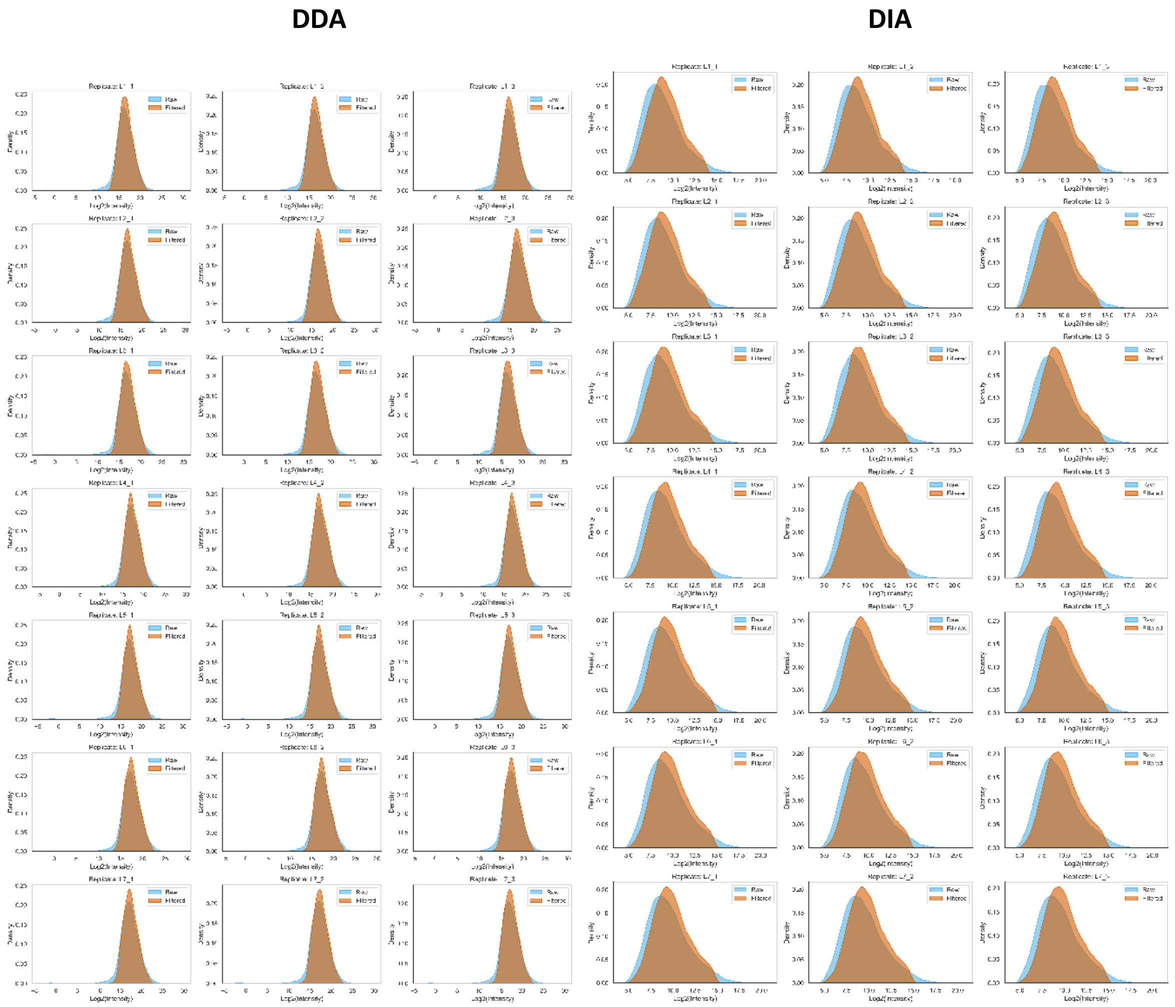


**Figure S1.** Kernel density estimates of $log_2$-transformed peptide intensities in DDA (left) and DIA (right) data before and after filtering, across all spike levels (L1–L7) and technical replicates (1–3). Each subplot corresponds to a specific replicate within a spike level. Raw intensities (blue) represent the unfiltered data, while filtered intensities (orange) include only peptides passing all quality control thresholds.

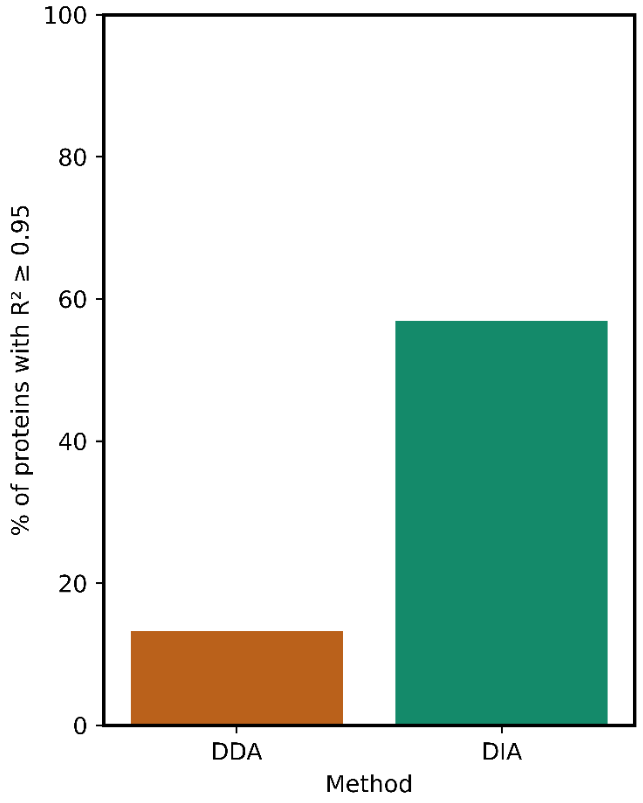


**Figure S2.** Percentage of proteins exhibiting high linearity ($R^2 \geq 0.95$) across spike levels in DDA and DIA workflows.

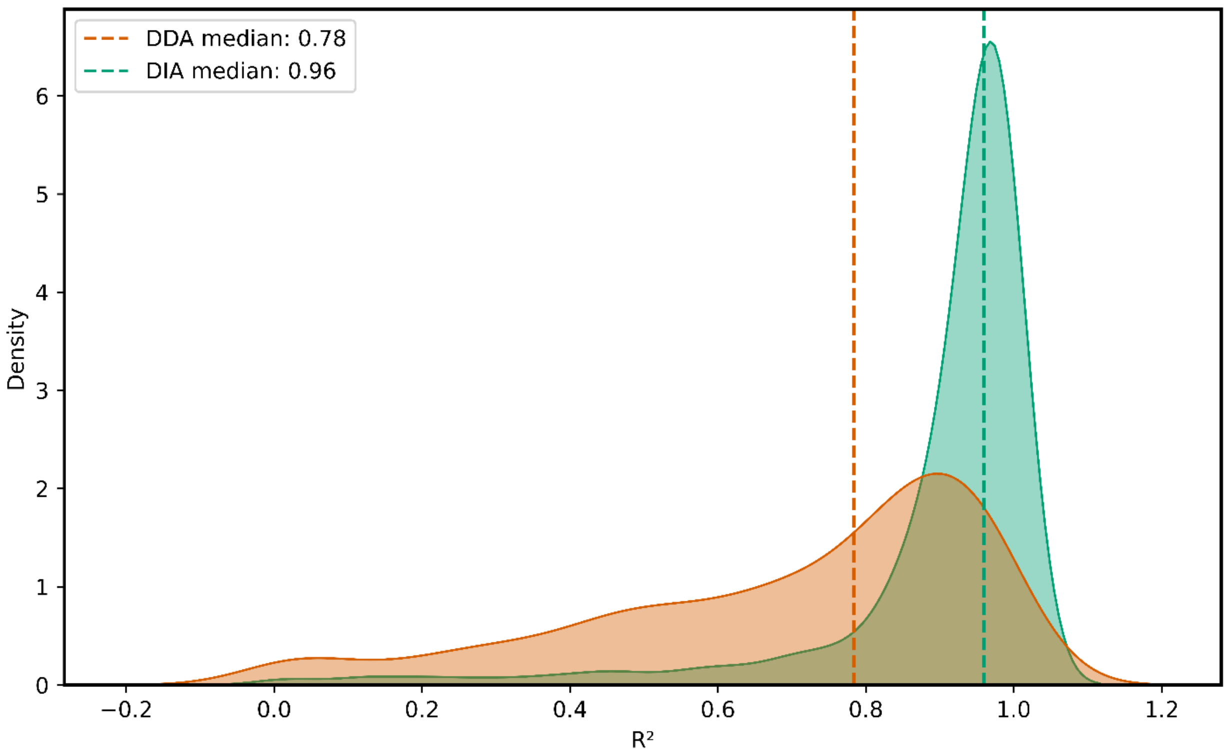


**Figure S3.** Kernel density distribution of protein-wise $R^2$ values from linear regression across spike levels, comparing DDA and DIA workflows. Vertical dashed lines denote group medians.
